# Limited significance of the in situ proximity ligation assay

**DOI:** 10.1101/411355

**Authors:** Azam Alsemarz, Paul Lasko, François Fagotto

## Abstract

In situ proximity ligation assay (isPLA) is an increasingly popular technique that aims at detecting the close proximity of two molecules in fixed samples using two primary antibodies. The maximal distance between the antibodies required for producing a signal is 40 nm, which is lower than optical resolution and approaches the macromolecular scale. Therefore, isPLA may provide refined positional information, and is commonly used as supporting evidence for direct or indirect protein-protein interaction. However, we show here that this method is inherently prone to false interpretations, yielding positive and seemingly ‘specific’ signals even for totally unrelated antigens. We discuss the difficulty to produce adequate specificity controls. We conclude that isPLA data should be considered with extreme caution.

## Introduction

A major challenge of cell biology is the ability to detect protein-protein interactions in situ. The current methods of choice are FRET and related techniques. However, this type of approach involves expression of tagged fusion proteins, has limited sensitivity, and often requires extensive optimization. The in situ proximity ligation assay (isPLA) has thus appeared as a very attractive, easy and ready to use alternative [1-10]. Its principle is based on the immunodetection of two antigens with a pair of primary antibodies raised in different species (Fig.1A). The two primary antibodies are then recognized by two species-specific secondary antibodies, called PLA probes, each linked to a unique short DNA strand. When the two PLA probes are in close proximity, the DNA strands can be used to recruit two additional connector oligonucleotides, which are ligated to form a DNA circle. This allows the synthesis of a single-stranded rolling circle PCR product, composed of hundreds of concatenated complements of the DNA circle, which is then visualized using a fluorescently-labelled complementary oligonucleotide probe. The maximal distance allowing this reaction is 40 nm, which is not quite small enough to demonstrate a physical interaction between the two antigens, but sufficient to support a very close ‘proximity’. It is certainly below the limit of optical resolution, thus potentially much more informative than classical fluorescence colocalization experiments. The potential applications of this method are huge, since it can be used in principle with any pair of antibodies, allowing co-detection of any endogenous antigen, including posttranslational modifications, such as specific phosphorylated sites. This flexibility explains its increasing popularity, in fields as diverse as cell biology, pharmacology, immunology, virology, proteomics, biomarkers for cancer, pathogen diagnostic, or even astrobiology [5,7,8,10-17].

**Figure 1.**
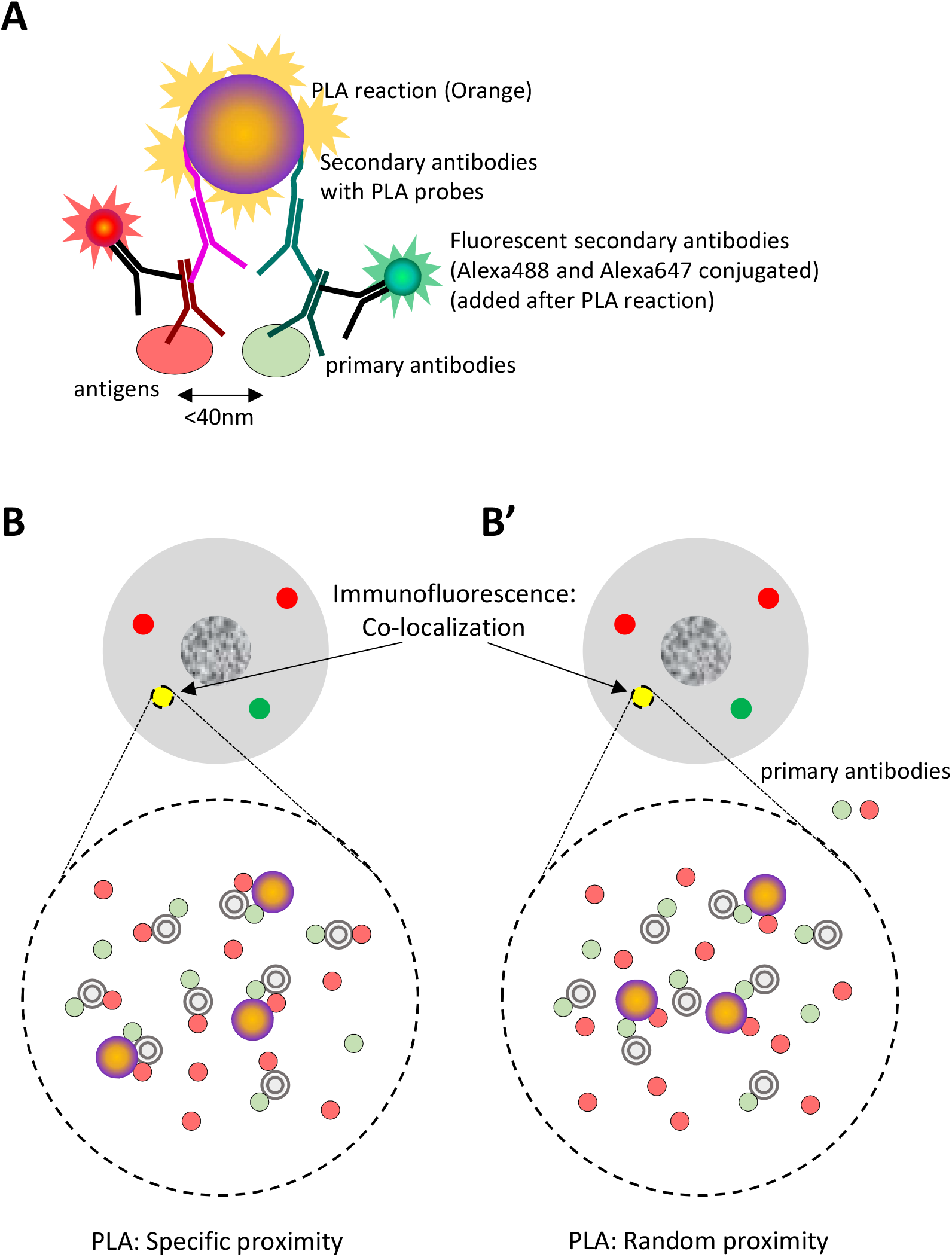
A. Principle of the in situ proximity ligation assay coupled to double immunofluorescence labelling. After incubation with two primary antibodies, secondary antibodies with PLA probes are added and the PLA reaction is performed (details of the reaction omitted). The reaction can only occur if the two antigens are closer than 40nm. In a subsequent step, regular fluorescently-labelled secondary antibodies are added in order to determine the distribution and levels of each primary antibody. In the experiments presented here, we used the orange fluorescent PLA reagent (emission at 579 nm), and green (Alexa488-conjugated) and far red (Alexa647-conjugated) secondary antibodies. Nuclei were counterstained with Hoechst (not shown). **B,B’. Specific versus fortuitous proximity.** Assuming that two antibodies yield a co-localization pattern (yellow spot in the cell drawing), we ask whether isPLA can further discriminate between a specific proximity due to association of the two antigens within a protein complex (or a subcellular structure, in the order of few tens of nanometers) or a fortuitous proximity due to the high antigen/antibody global concentrations within the region observed by immunofluorescence. In B’, the green antibody marks a protein associated with the subcellular structure, the red antibody recognizes a randomly distributed antigen. Antigens, secondary antibodies and probes are omitted for clarity sake.

Unfortunately, when attempting to apply this approach, we were surprised to obtain robust positive results for pairs of antigens which, although partially yielding overlapping immunofluorescence signals, could not possibly establish any direct or indirect interaction. A closer consideration of the principle of isPLA suggested the possibility that indeed a positive signal may be generated for any pair of antigens, provided that a subset of the primary antibodies would happen to bind sufficiently close to each other. Yet, the goal of a proximity assay should be to reveal the presence of two antigens within structures in the tens of nanometer range (a macromolecular complex, a vesicle, or a membrane subdomain), and to discriminate between these relevant cases and a mere random proximity, for instance the close but fortuitous encounter of a plasma membrane protein and an unrelated soluble cytosolic protein (Fig.1B).

We therefore evaluated the capacity of isPLA to differentiate between an actual interaction *versus* random proximity (Fig.2B,B’). We unambiguously conclude that isPLA yields positive signals in a variety of conditions that are irrelevant both in terms of protein-protein interactions or refined subcellular localization. In fact, attempts to set adequate specificity controls as performed in our study, and which would be necessary to support the existence of a biologically relevant proximity, were, to our knowledge, not implemented in other studies. As currently performed, isPLA is most likely to yield a large number of false positives and to lead to unsupported conclusions.

**Figure 2.**
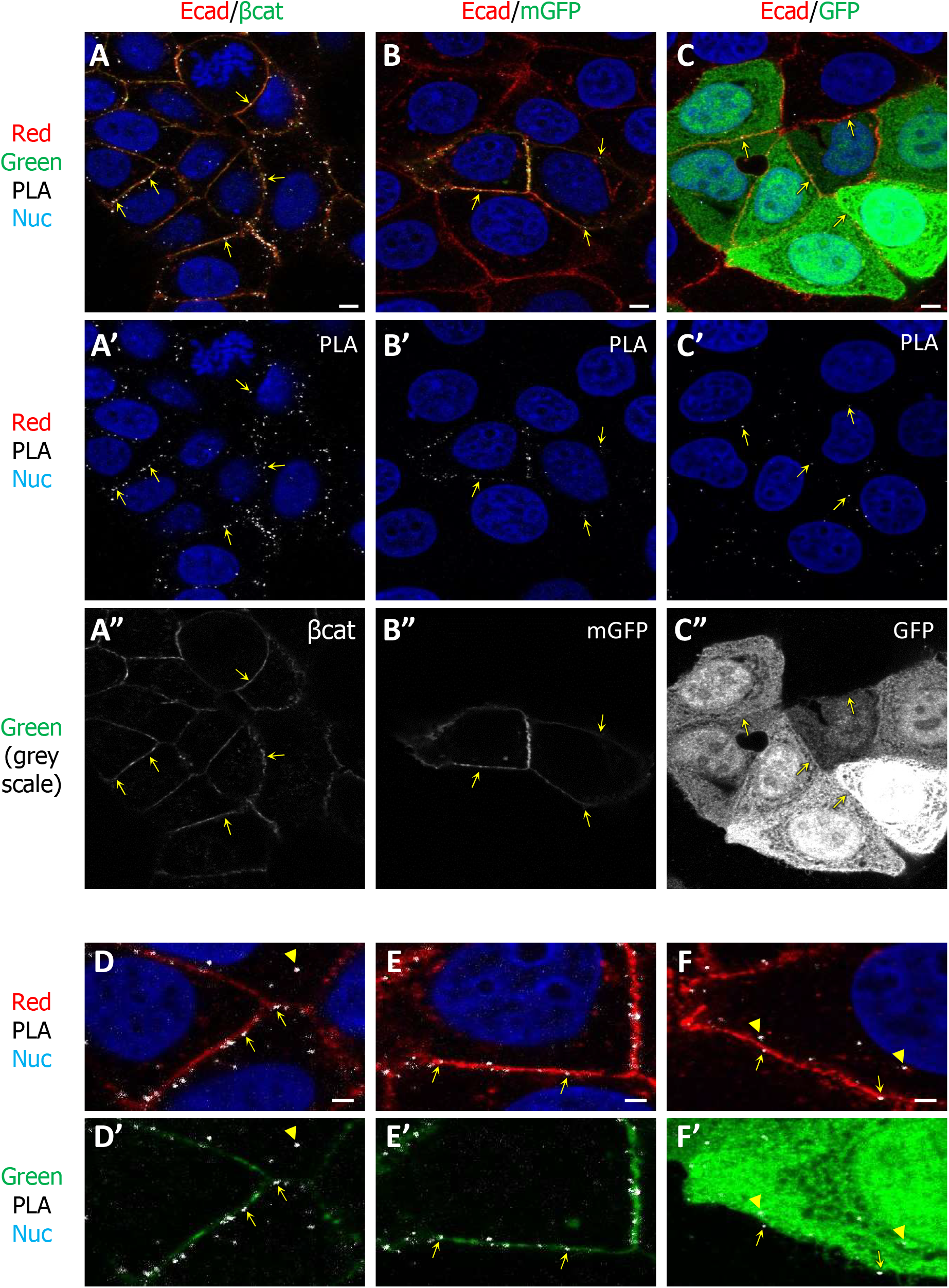
isPLA signal at the plasma membrane. **A.** isPLA for endogenous E-cadherin and β-catenin. **B, C.** isPLA using anti-E-cadherin and anti-GFP antibodies on cells ectopically expressing membrane-targeted GFP (mGFP)(B) or soluble GFP (GFP)(C). In all cases, positive isPLA signal was observed along the plasma membrane (arrows). **D-F.** High magnification from A-C. Arrows point to spots along the plasma membrane. Arrowheads in F point to examples of PLA corresponding to cytoplasmic, E-cadherin-positive structures. See supplemental Figure S1 for additional examples, negative controls, and quantifications. All images were captured by laser scanning confocal microscopy using the exact same settings. Scale bars, A-C, 5μm; D-F, 2μm.

## Results

In the series of tests presented here, we set to evaluate the occurrence of PLA positive signals for different situations, comparing pairs of antibodies against known interacting proteins and pairs recognizing unrelated antigens. The PLA results were analysed both in terms of distribution of the fluorescent spots and of spot density. An additional classical double immunofluorescence labelling was performed on the same samples (Fig.2A), which allowed to directly verify the distribution of the two primary antibodies and compare relative intensities between conditions. The standard negative control for isPLA is the omission of one of the primary antibodies. In our hands, such control always yielded little or no PLA reaction, but as discussed below, this cannot be considered as an acceptable control.

In this survey, we tested PLA for well-established residents of three cellular structures, the plasma membrane (E-cadherin), cytoskeleton (α-tubulin) and nucleus (transcription factor Sox9). We used ectopically expressed soluble GFP as randomly distributed protein. As a general marker for the plasma membrane we expressed mGFP, i.e. GFP fused to a sequence that becomes palmitoylated and thus efficiently anchored to the inner leaflet of the plasma membrane. GFP and mGFP were detected with a specific anti-GFP antibody. Note that using ectopic GFP enabled us to monitor isPLA for a wide range of expression levels and immunofluorescence intensities, which could be compared to signals obtained after labelling endogenous proteins.

We also included in our tests non-specific IgGs. These are normally used as classical negative control in traditional immunofluorescence, but they are also well known to broadly label the cytoplasm and the nucleus, to an extent that depends on the concentration used. We thus used them here as an example of widespread non-specific antibody labelling. Upon adequate titration, they yielded an immunofluorescence signal in the range obtained with the other antibodies (Figs.3, S2 and S3). We stress here this important distinction between controls for <u>specific immunolabelling</u> or for <u>specific proximity</u>. For the latter purpose, any type of negative control, which removes binding of one of the primary antibodies (e.g. omitting the antibody, knocking out the antigen, competing with the corresponding peptide) will give trivially a low or blank signal, but will give not information about the specificity of the positive reaction. Instead, one needs to be able to evaluate side by side isPLA produced by a candidate pair of interactors, as depicted in Fig.1B, or produced non-specifically by random adjacent antibodies present in the region of interest at similar levels, as depicted in Fig.1B’. To our knowledge, this type of control has never been implemented for isPLA.

We compared isPLA between E-cadherin and its direct cytoplasmic interactor β-catenin, E-cadherin and mGFP, or E-cadherin and soluble GFP (Fig.2 and suppl. Fig.S1),. The anti-E-cadherin antibody recognized an epitope on the extracellular domain. We found that the plasma membrane was decorated with PLA signal in all three cases (Fig.2, arrows). The density of PLA spots appeared to be roughly proportional to the intensity of the β-catenin/GFP immunofluorescence signals, thus to the density of the primary antibodies. This relationship was quantified through line scans along the membrane (Fig.S1H-I), and the data were plotted as PLA density as function of green fluorescence intensity signal, representing the relative levels of anti-β-catenin or anti-GFP antibodies (Fig.S1J): While the frequency of PLA spots tended to be on average slightly higher for the E-cadherin-β-catenin pair, and lowest for the E-cadherin-GFP pair, the three distributions largely overlapped (enlargement in Fig.S1J). We concluded that for similar levels of primary antibodies, PLA could not effectively discriminate between the specific E-cadherin-β-catenin interaction and the other conditions.

We then tested isPLA for tubulin and soluble GFP or mGFP, or non-specific IgGs (Fig.3 and Fig.S2). Robust PLA was obtained in all three cases. The spots decorated microtubules in the first and last conditions (Fig.3A,C), consistent with the widespread distribution of the anti-GFP and non-specific IgG signals. For the tubulin-mGFP pair, they concentrated along the plasma membrane, where the two antibodies mostly overlapped (Fig.3B). Again, the density of PLA spots was related to the global antibody levels (e.g. compare cells 1 and 2, in Figure 3A, which express different GFP levels), although in detail the PLA position did not necessarily correlate with sites of highest immunofluorescence signal. Standard negative controls, i.e. cells not expressing GFP (Fig.3A,B and Fig.S2A,B), or omission of the anti-tubulin antibody (Fig.S2D) gave little to no PLA signal.

**Figure 3.**
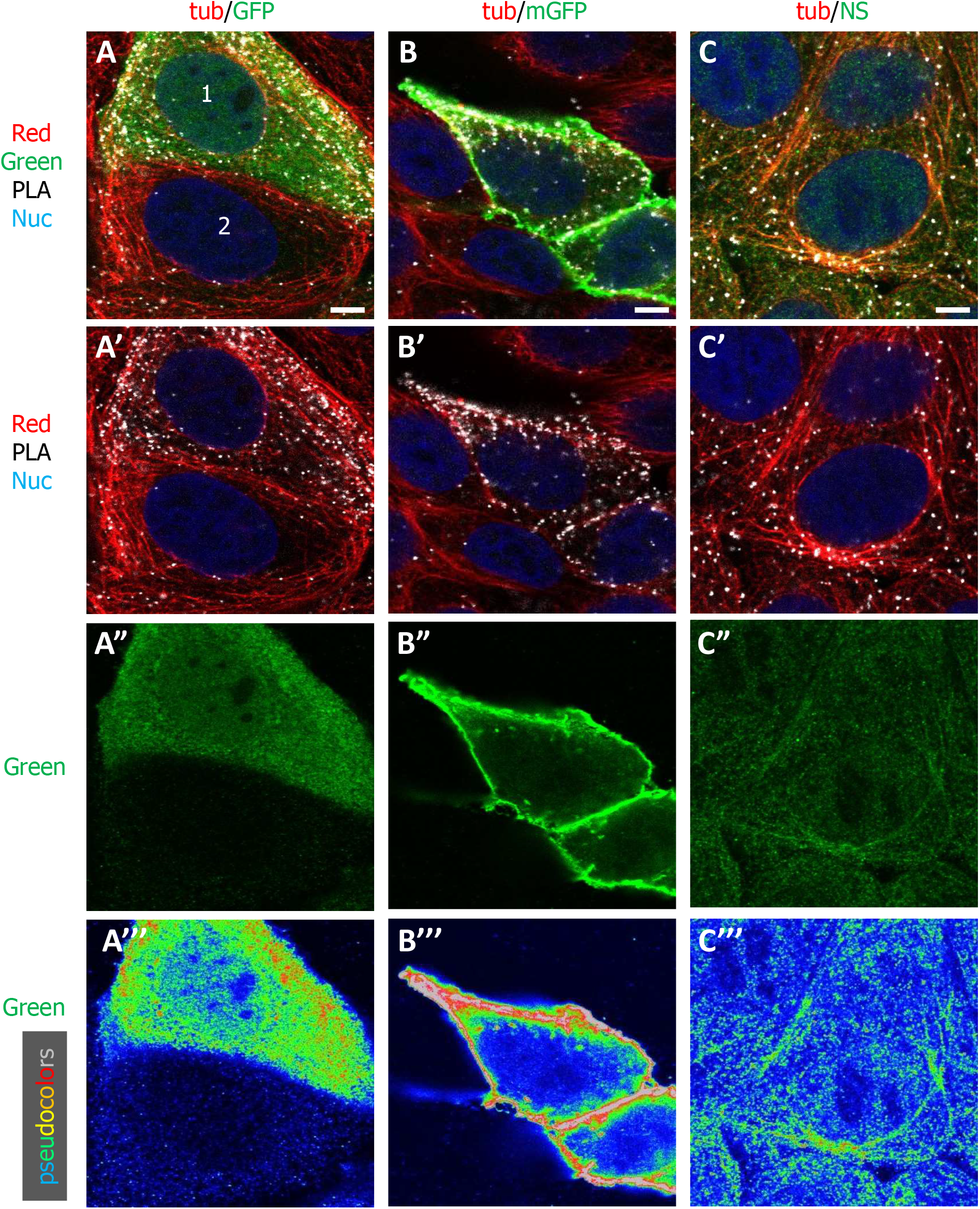
isPLA signal at microtubules. **A.** isPLA for endogenous α-tubulin and soluble GFP. Detail of two cells expressing moderate (cell 1) and weak levels (cell 2) of soluble GFP (pseudocolors included for better comparison, see Fig.S2 for full fields). Numerous spots are observed along microtubules. Spots are denser in the cell expressing higher levels of GFP. **B.** isPLA for endogenous α-tubulin and membrane-targeted GFP (mGFP). Detail of two cells expressing mGFP. Numerous spots are found at the cell periphery, where most of the tubulin and mGFP signals overlap. **C.** isPLA using anti-α-tubulin and a non-specific rabbit serum. The dilution of the serum was adjusted as to yield a non-specific signal of an intensity comparable to the intensity for the anti-GFP antibody. Microtubules are decorated by PLA spots. Scale bars, 5μm.

Finally, we compared isPLA for antibodies raised against Sox9 and Sam68, a nuclear factor recently shown to physically interact with Sox9 [18] (Fig.4A), or for anti-Sox9 and a non-specific antibody (Fig.4B). An additional interest of the Sox9-Sam68 interaction was its proposed enrichment in the peripheral region of the nucleus, based on isPLA [18]. We reproduced this distribution (Fig.4A,A’, histogram in panel C), which was in stark contrast with the immunofluorescence staining indicating a relatively homogenous distribution of the primary antibodies (Fig.4A”, quantification in Fig.S3F). Strikingly, however, a very similar pattern was observed when the anti-Sam68 antibody was replaced with non-specific IgGs (Fig.4B,B’,D), indicating that it did not represent a specific sub-nuclear site of interaction (discussed below). Note that PLA spots were also abundant in the cytoplasm signal, which was surprising considering the rather low cytoplasmic immunofluorescence signals for Sox9 and Sam68 (see discussion). Negative controls omitting anti-Sox9 or anti-Sam68 antibodies were blank for PLA (suppl. Fig.3D,E).

**Figure 4.**
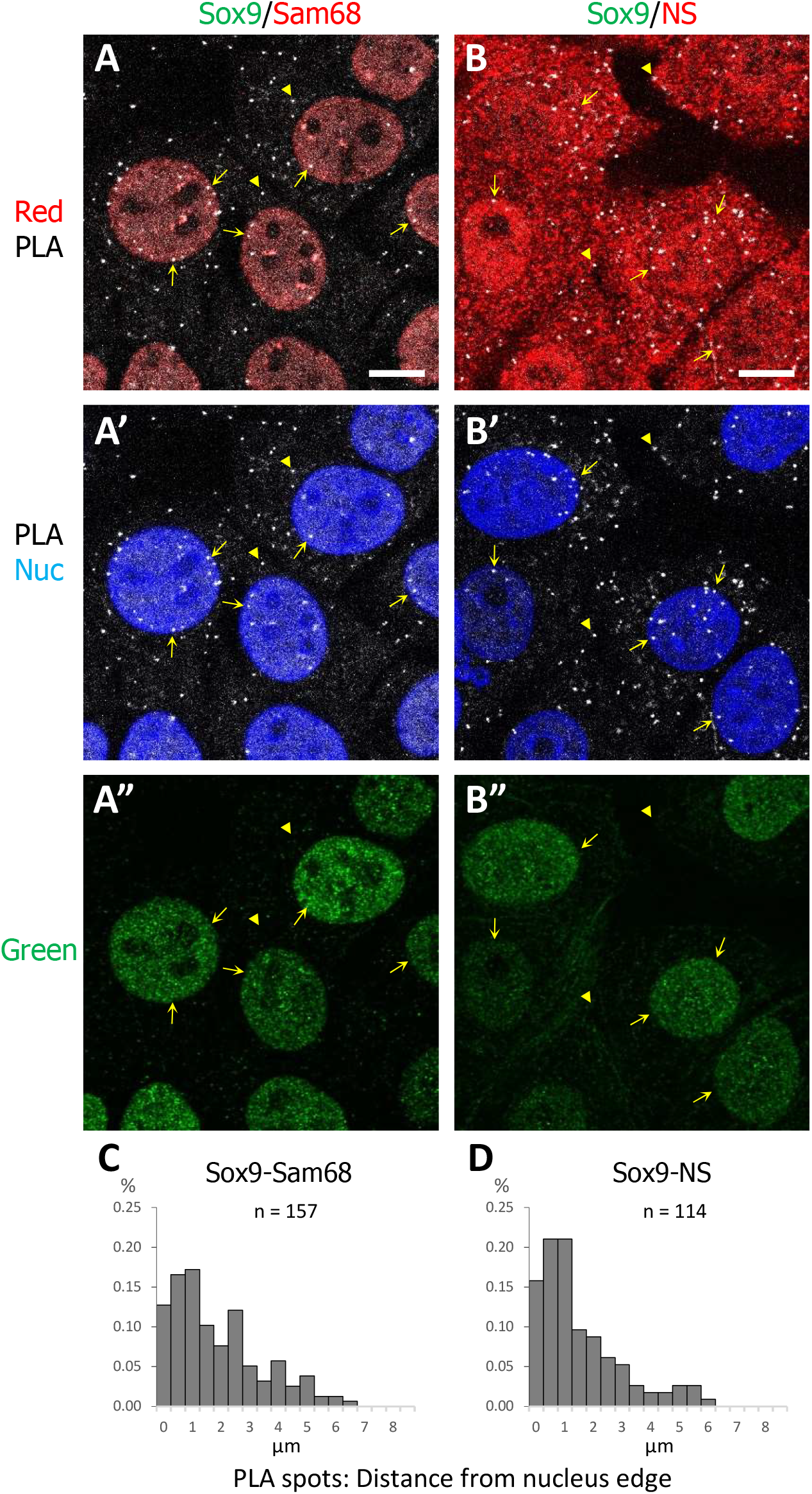
isPLA in the nucleus. **A,B.** isPLA for Sox9, chosen as example of nuclear protein, and either Sam68, a candidate nuclear interacting partner (A), or non-specific serum (NI)(B). For both conditions, PLA spots were observed both in the nucleus (arrows) and in the cytoplasm (arrowheads). Nuclear spots are mostly found at the periphery of the nuclei (see Fig.S3 for more details). **C.** Negative control, anti-Sox9 antibody omitted. Scale bars, 5μm.

## Discussion

The series of tests presented here show that the proximity ligation assay can produce positive reactions under a variety of situations, including conditions that bear little to no significance in terms of actual specific proximity between macromolecules (Fig.5). This is a critical issue, which, retrospectively, appears inherent with the principle of the assay: Because the annealing of the probes requires the right distance and positioning of two antibodies, as well as the contribution of multiple reagents, the PLA reaction is intrinsically stochastic, with a probability increasing with, among other factors, the local density of the antibodies. It has been assumed that such high density reflects a specific concentration of two antigens, resulting from their physical interaction or from their localization to the same subcellular structure. These are certainly favourable conditions for PLA, but the reaction may also be generated with any pair of antigens, even if they clearly do not interact, as shown here for GFP and cadherin or tubulin, the only apparent condition being a partial overlap of their distribution (Fig.1B’ and Fig.5). The resulting PLA pattern will primarily be determined by this overlap, rather than by the actual distribution of the antigen, and the density of spots generated in these overlapping regions will depend directly on the density of bound antibodies, not necessarily on a specific local accumulation of the two antigens. This easily explains the striking, seemingly “specific”, PLA decoration of membranes or microtubules obtained in our tests that used broadly distributed antigens. The illusion of a “specific proximity” may appear particularly convincing in cases where the two antibodies would appear by immunofluorescence to mark preferentially the same discrete structure (e.g. centrosome, cilium, or nucleolus, see symbolized yellow circle in Fig.1B,B’). These considerations lead us to conclude that in its current form, isPLA does not bring more information than classical immunofluorescence co-localization.

**Figure 5.**
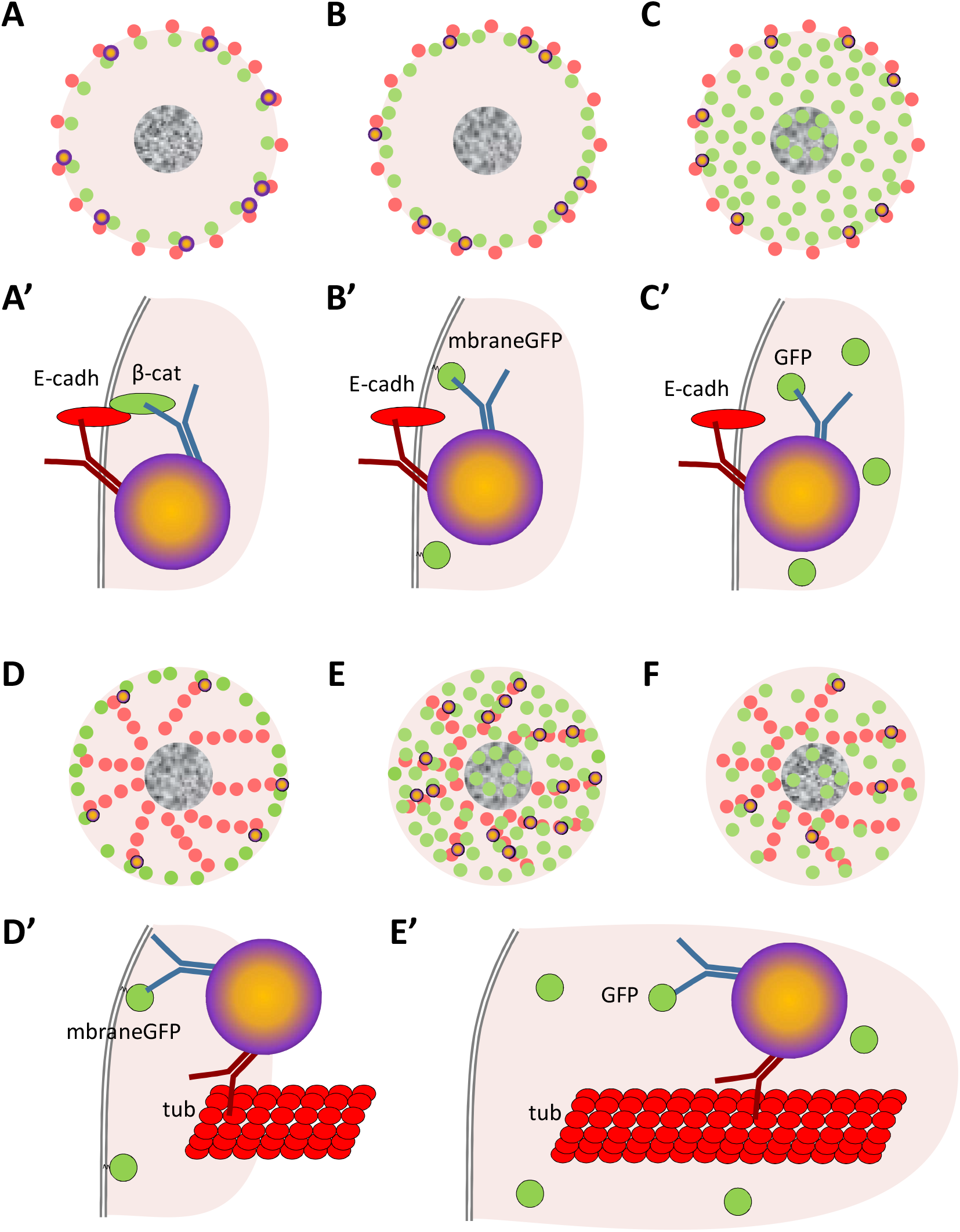
Summary diagram. Panels A-F show the general cellular distribution of the PLA signal (orange dark-circled dots), and of the primary antibodies (green and red). Panels A’-E’ represent the detailed situation generating PLA in each case. Secondary antibodies and probes are omitted for clarity’s sake. isPLA yields positive signals not only for biologically meaningful pairs of antigens, here E-cadherin and β-catenin (A), but potentially for any pair of antigens, provided that a small fraction of the two primary antibodies bind sufficiently close to allow for the ligation reaction (B-F). The pattern thus obtained may then give the illusion of a “specific interaction”, for example along the plasma membrane (B,C), at the intersection between microtubules and the plasma membrane (D), or decorating microtubules (E,F), even when one of the targeted antigen is widely distributed, such as in the case of soluble GFP (C,E,F). The density of the PLA spots depends on the levels of primary antibodies (E and F), which is dictated by antigen abundance, antibody concentrations, and additional parameters (see discussion).

To be able to extract more useful information from isPLA, it would be imperative to set controls and criteria that would convincingly define a meaningful “proximity”. As mentioned above, classical controls only verify the specificity of the antibody, not of the proximity reaction. The standard negative control of the isPLA method, i.e. simple omission of one of the two primary antibodies [6,9,16], is insufficient, as it only controls for the potential non-specific binding of the secondary antibodies (and here PLA probes), which is generally very low anyway. We have confirmed here that indeed little to no PLA reaction is observed under this condition. Another previously suggested control is the use of a sample missing one of the antigens, either naturally or through knock-out/knock-down [6,9], but again, this control can only validate the specificity of the antibodies, not of the isPLA.

One clearly needs much more stringent tests. One possibility would be to compare the density of PLA spots to the levels of primary antibodies localizing at the structure of interest, taking as reference point a “negative control” condition (suppl. Fig.S1). Unfortunately, we readily noticed serious obstacles to implementing such a control. A major problem is that the assay relies on antibody binding, which, unlike for instance FRET, does not necessarily faithfully reflect the position and levels of the molecules under investigations. Indeed, antibody binding varies widely, depending on affinity, sensitivity to fixation, and antigen availability/masking. In practice, we do not believe there is any objective criterion that would allow us to set appropriate antibody dilutions in order to quantitatively compare “positive” and “negative control” signals.

Our tests reveal additional levels of complexity and highlight the difficulty in drawing conclusions from PLA experiments. We were surprised that PLA for E-cadherin-β-catenin, which form 1:1 complexes, was only marginally more effective than PLA for the “random” E-cadherin-mGFP pair (Fig.2 and Fig.S1). It is quite easy to conceive that despite the known direct interaction of E-cadherin with β-catenin, the two epitopes targeted by the antibodies may not be in the optimal configuration for efficient PLA. Note indeed that in addition to the absolute distance between the epitopes, their relative orientation (and therefore the position of the antibodies) may also influence the probability of a positive PLA reaction. Partial antigen masking within the dense adhesive structures could also decrease the probability of simultaneous binding of the two antibodies to the same cadherin-catenin complex. In any case, this example clearly shows that isPLA using a well-characterized pair of interacting partners may not be as efficient as originally expected.

The influence of complex parameters such as accessibility and orientation is supported by the line scans presented in supplemental Fig.S1, which show that PLA spots do not necessarily coincide with local peaks of antibody concentration. This is a general observation, which we made for all conditions tested (not shown). Furthermore, the high frequency of cytoplasmic PLA spots in the case of Sox9-Sam68 (Fig.4), despite the strong accumulation of both antibodies in the nucleus also argues in favour of this hypothesis.

Our analysis of nuclear PLA raises an additional issue. PLA enrichment at the periphery of this organelle, initially reported for the Sox9-Sam68 pair, was also observed for the non-specific anti-Sox9/non-specific IgG pair (Fig.4 and suppl. Fig.S3), which suggests the existence of an intrinsic bias independent of specific protein-protein interactions. We think that it may be linked to the density of the nuclear content, which, although not sufficient to significantly restrict the diffusion of primary and secondary antibodies (Fig.S3G), could be a more serious obstacle for the PLA reaction, which requires that the simultaneous convergence of multiple components (oligonucleotides, ligase, polymerase, fluorescent probes) on the same spot. While the nucleus is arguably the densest and largest structure of the cell, similar considerations about limiting diffusion and accessibility may apply in subtler ways to other parts of the cell.

In summary, we conclude that in its present form, isPLA cannot be trusted as source of information about localization/interaction at the subcellular level.

## Material and methods

### Cells and transfections

Human breast cancer MCF7 cells (ATCC HTB-22) were grown in Dulbecco’s Modified Eagle’s medium (Life technologies, 31966047) with 10% FBS (Life technologies, 10500064). Cells were transfected with pCS2-eGFP or pCS2-mGFP [19] using jetPRIME transfection reagent (Ozyme, POL114-07), according to manufacturer instructions. For imaging, MCF7 cells were seeded on 12 mm No. 1.5 coverslips coated with 50ug/ml collagen type I (Corning, 354236).

### Antibodies

Antibodies:
| Antigen | Species | Conc.<br>(μg/ml) | Supplier | Cat. number |
| --- | --- | --- | --- | --- |
| GFP | Mouse | 1.25 | Life technologies | A11120 |
| E-cadherin | Rabbit | 0.33 | Cell Signaling | 3195S |
| β-catenin | Mouse | 0.83 | Invitrogen | 13-8400 |
| α-tubulin | Mouse | 2.5 | Sigma | T6199 |
| non-specific IgG | Rabbit | 20 | Santa Cruz Biothechnology | SC-2027 |
| SOX9 | Mouse | 1.0 | Sigma | 1.0 |
| Sam68 | Rabbit | 4 | Santa Cruz Biothechnology | SC-333 |

### Proximity Ligation Assay and Immunofluorescence

Cells were fixed for 10 min with 3.7% paraformaldehyde (EMS #15714) in PHEM buffer (60 mM Pipes, 25 mM Hepes, 8 mM EGTA 4 mM MgCl2), followed by 10 min permeabilization with 1% Triton X100 (Applichem. Panreac., A4975) in phosphate buffer saline, and blocking with 20% sheep serum in phosphate buffer saline for 1h. Primary antibodies in blocking buffer serum for 2h were added.

Proximity ligation assay was performed using the Duolink kit (Sigma-Aldrich DUO92102), according to manufacturer’s protocol, using Orange red reagent (excitation 554 nm, emission 579 nm). After completion of the protocol, two additional secondary antibodies, coupled to green (Alexa488, ThermoFisher) and to far red (Alexa 647, ThermoFisher) dyes, were added at 5μg/ml, with the goal to directly detect each of the two primary antibodies. Samples were finally counterstained with Hoechst 33258 (5 μg/ml, Invitrogen, 33342). Images of the four colour signals were collected using a Leica SP5 laser scanning confocal microscope, with a 63X NA objective.

## Acknowledgments

We want to thank David Rozema and Dominique Hemlinger for critical reading of the manuscript, and Peggy Raynaud for generous gift of antibodies. F.F. research is supported by the ANR grant ANR-14-ACHN-0004–ICM and by the Labex EpiGenMed.

---

**Supplemental Figure S1. isPLA signal at the plasma membrane.**

**A-E.** Comparison of isPLA for endogenous E-cadherin with β-catenin and with different levels of ectopic mGFP/GFP. A,B and D are as in Figure 1.

**F,G.** Negative controls (omission of anti E-cadherin, or anti-β-catenin primary antibodies).

**H,I.** Examples of line scans along plasma membranes. Plots represent red (anti-E-cadherin) and green (anti-β-catenin or anti-GFP) fluorescence intensities. Arrows indicate the position of PLA spots.

**J.** PLA spot densities along plasma membranes plotted as function of green fluorescence intensity.

**Supplemental Figure S2. isPLA signal along microtubules.** A-D. Merge of the four channels. A’-D’. Merge of tubulin, PLA and Hoechst signals. A”-D”. PLA signal alone. A’’’-D’’’. Green (GFP or non-specific) signals displayed as pseudocolors.

**A,B.** isPLA for α-tubulin and GFP (A,B). C. isPLA for α-tubulin and non-immune serum (C).

**D.** Negative control (omission of anti-GFP antibody). B and C are full fields of images of Fig.2.

**Supplemental Figure S3. isPLA signal in the nucleus.**

**A,B.** Two examples of siPLA Sox9-Sam68. A correspond to the full field of A in Figure 3. **C.** siPLA Sox9-non-sepcific serum (NS). Full field of B in Figure 3. D. Negative control (omission of anti-Sam68).

**E,F.** Histogram of distribution of PLA spots in nuclei, measured in μm from the edge of the nucleus. Median values are 1.1 μm for Sox9-Sam58 and 0.8μm for Sox9-NI. Average nucleus diameter was ~ 8 μm.

**G.** Superposition of PLA spot distribution (as in E) with average fluorescence intensity profiles for Hoechst, anti-Sox9 and anti-Sam68.

